# Hippocampus, retrosplenial and parahippocampal cortices encode multi-compartment 3D space in a hierarchical manner

**DOI:** 10.1101/200576

**Authors:** Misun Kim, Eleanor A. Maguire

## Abstract

Humans commonly operate within 3D environments such as multi-floor buildings and yet there is a surprising dearth of studies that have examined how these spaces are represented in the brain. Here we had participants learn the locations of paintings within a virtual multi-level gallery building and then used behavioural tests and fMRI repetition suppression analyses to investigate how this 3D multi-compartment space was represented, and whether there was a bias in encoding vertical and horizontal information. We found faster response times for within-room egocentric spatial judgments and behavioural priming effects of visiting the same room, providing evidence for a compartmentalised representation of space. At the neural level, we observed a hierarchical encoding of 3D spatial information, with left anterior hippocampus representing local information within a room, while retrosplenial cortex, parahippocampal cortex and posterior hippocampus represented room information within the wider building. Of note, both our behavioural and neural findings showed that vertical and horizontal location information was similarly encoded, suggesting an isotropic representation of 3D space even in the context of a multi-compartment environment. These findings provide much-needed information about how the human brain supports spatial memory and navigation in buildings with numerous levels and rooms.

## Introduction

We live in a three-dimensional (3D) world. The ideal method to enable navigation in 3D space would be to have a 3D compass or global positioning system (GPS) that identifies direction and distance in relation to all 3 axes in an isotropic manner. Such a 3D compass system would seem to be essential for animals who fly or swim. Indeed, place cells and head direction cells found in flying bats were observed to be sensitive to all three axes (Yartsev and Ulanovsky 2013; Finkelstein et al. 2014). Behavioural experiments with fish also indicated a volumetric 3D representation of space (Burt de Perera et al. 2016). However, as suggested by Jeffery et al. (2015), extending spatial encoding from 2D to 3D space comes with complications such as the non-commutative property of 3D rotation, and a fully volumetric representation of 3D space might be costly and unnecessary for certain environments and species. When an animal’s movement is restricted on the earth’s surface due to gravity, its position can be identified by two coordinates on that surface and a quasi-planar representation could be more efficient than a volumetric 3D representation.

Multi-floor buildings are the most common type of working and living spaces for humans today. Regionalisation is a key characteristic of these environments - multiple floors stacked on top of each other and multiple rooms located side by side on a floor. When we navigate within multi-floor buildings, we can use hierarchical planning rather than using a 3D vector shortcut or volumetric 3D map. For example, we decide which floor to go to (“second floor”), and which room on that floor (“the first room nearest the stairs”), then the location within the room (“the inside left corner of the room”). The regionalisation and hierarchical representation of space involving multiple scales has been consistently observed (Hirtle and Jonides 1985; Han and Becker 2014; Balaguer et al. 2016), but it is not fully understood how spatial information about multiple scales is encoded at the neural level, particularly in a 3D context.

One obvious question is whether a common neural representation is used for each compartment (room). Using a generalised code to register local information is an efficient strategy compared to assigning unique codes for every location in an entire environment in the context of repeating substructures. Moreover, a common local representation can be seen as a “spatial schema” that captures the essence of an environment and helps future learning of relevant environments or events (Tse et al. 2007; Marchette et al. 2017). The retrosplenial cortex and hippocampus are candidate brain regions for the encoding of within-compartment, local information. Place cells in the hippocampus are known to repeat their firing fields in a multi-compartment environment (Derdikman et al. 2009; Spiers et al. 2015). Moreover, human fMRI studies have shown that the hippocampus contains order information that generalises across different temporal sequences (Hsieh et al. 2014), and the retrosplenial cortex contains location codes that generalise across different virtual buildings (Marchette et al. 2014).

Another question is where in the brain each compartment is represented within the larger environment in order to complement the local room information. To the best of our knowledge, no study has simultaneously interrogated the neural representation of local spatial representations and the compartment information itself. The hippocampus might contain both types of information. It has been suggested that the hippocampus represents spatial information of multiple scales down its long axis. For example, the size of place fields is larger in ventral hippocampus than dorsal hippocampus in rats (Kjelstrup et al. 2008). In a human fMRI study, increased activation in posterior (dorsal) hippocampus was associated with a fine-grained spatial map, whereas the anterior (ventral) hippocampus was linked with coarse-grained encoding (Evensmoen et al. 2015).

It is also interesting to consider the question of whether vertical and horizontal information is equally well encoded in a 3D environment. In other words, is it the case that when a room is located directly above another room, are they as equally distinguishable as two rooms that are side by side on the same floor? At the neural level in rats, place cells and grid cells show vertically elongated firing fields, implying reduced encoding of vertical information, and it has been proposed that this is evidence of the quasi-planar representation of 3D space (Hayman et al. 2011; Jeffery et al. 2013). However, this asymmetry could have arisen from the repeating shape of the environment. By contrast, Kim et al. (2017) found that the human hippocampus encoded vertical and horizontal location information equally well in a 3D virtual grid-like environment during fMRI scanning.

Behaviourally, there are mixed results in the literature in relation to vertical-horizontal symmetry/asymmetry. In a study by Grobéty and Schenk (1992), rats located the vertical coordinate of a goal earlier than the horizontal coordinate, whereas in Jovalekic et al. (2011), rats prioritised horizontal movements and foraged at the horizontal level before moving to the next level. In humans, a group who learned the location of objects in a virtual multi-floor building along a floor route had, overall, better spatial memory than a group who learned along a vertical columnar route, suggesting a bias towards the floor-base representation (Thibault et al. 2013). However, another study reported that twice as many participants reported a columnar representation of a building than a floor representation (Büchner et al. 2007).

Considering active navigation in a real buildings, Zwergal et al. (2016) found that participants were better during horizontal navigation than navigation across multiple floors. However, this difference could have arisen from reduced attention to visual landmarks during vertical navigation. It has also been reported that dogs correctly remembered horizontal locations within a floor but not the vertical floor itself (Brandt and Dieterich 2013). However, it is not clear whether the dogs’ navigation errors were due to inherent differences in vertical and horizontal encoding, or because of unequal landmark information consequent upon a lack of visual and odour controls in this study. Hummingbirds have been reported to be more accurate at vertical locations in a cubic maze, while the opposite pattern was observed for rats (Flores-Abreu et al. 2014).

We sought to address the issues outlined above in order to provide much-needed information about how regionalisation of space is realised at the neural level, in particular in a 3D context. Participants learned the locations of paintings in a virtual multi-floor gallery building. Before scanning, we compared their spatial judgments within and across vertical and horizontal boundaries. Then participants performed an object-location memory test while being passively moved in the virtual building during fMRI scanning. Repetition suppression analysis was used to ascertain which brain regions represented the local information within a room, or the room information within the building. In addition, we also asked whether vertical and horizontal room information in the brain was symmetrically or asymmetrically represented.

## Materials and Methods

### Participants

Thirty healthy adults took part in the experiment (15 females; age 23.7±4.6 years; range 18-35 years; all right-handed). All had normal or corrected to normal vision and gave informed written consent to participation in accordance with the local research ethics committee.

### The virtual environment

The virtual environment was a gallery building. There were 4 identical-looking rooms within the building, two rooms on each of two main floors (Fig. 1A, C). There was a unique painting located in each of the four corners of a room, resulting in 16 unique locations in the building. The paintings were simple and depicted animals or plants such as a dog, rose or koala bear. Painting locations were randomised across the participants, therefore spatial location was orthogonal to the content of the painting associated with it. The virtual environment was implemented using Unity 4.6 (Unity Technologies, CA, USA). A first-person-perspective was used and the field-of-view was ±30° for vertical axes and ±37.6° for horizontal axes. During the pre-scan training, the stimuli were rendered on a standard PC (Dell Optiplex 980, integrated graphic chipset) and presented on a 20.1 inch LCD monitor (Dell 2007FP). The stimuli filled 70% of the screen width. The same PC was used during scanning, and the stimuli were projected (using an Epson EH-TW5900 projector; resolution 1024 x 768) on a screen at the back of the MRI scanner bore and participants saw the screen through a mirror attached to the head coil.

**Figure 1.**
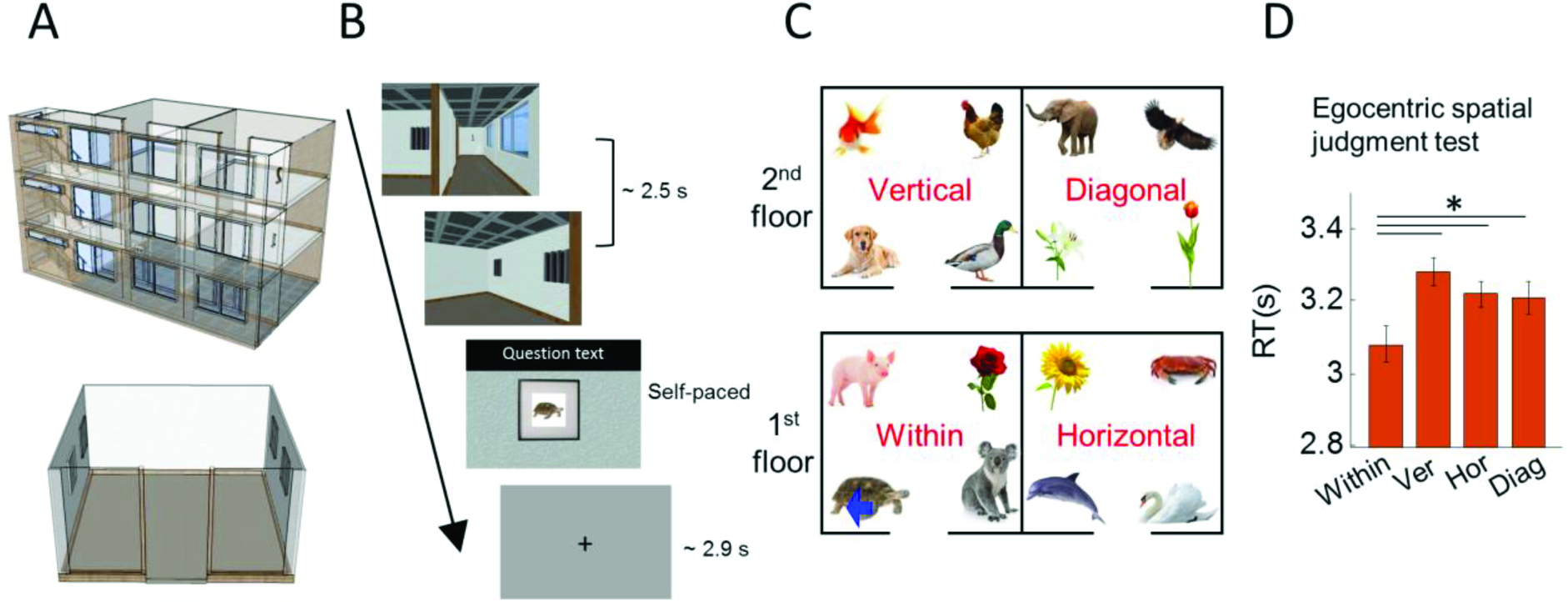
Stimuli and experimental design. (A) Top panel, overview of the virtual gallery building with transparent walls for display purposes. Bottom panel, overview of one room with transparent walls for display purposes. (B) On each trial, participants were virtually transported to one of the paintings from a corridor, and then the participant performed a spatial memory task. During the pre-scan egocentric judgments test, they were asked to make spatial judgments about the locations of other paintings, e.g. “Is the pig on your left?” (mean RT 3.2 s). During the scanning test, they were asked to indicate whether the painting was the correct one or not for that location, “Is this picture correct?” (mean RT 1.3 s). (C) An example layout of 16 paintings located in the 4 rooms of the gallery. The within, vertical, horizontal and diagonal rooms were defined relative to a participant’s current location. In this example, the participant was standing in front of the turtle painting (blue arrow). (D) Spatial judgments were significantly faster for the within-room (Within) condition. There were no differences between vertical (Ver), horizontal (Hor) or diagonal (Diag) rooms. Error bars are SEM adjusted for a within-subjects design (Morey 2008). *p<0.05.

### Tasks and procedure

Each participant completed the tasks in following order: learning prior to scanning, a pre-scan egocentric judgment task and the object location memory task during scanning (which was preceded by a short practice of the scanner task).

#### Learning prior to scanning

We allocated 20 minutes for the initial free exploration learning phase, but allowed participants to proceed to the test phase before 20 minutes had elapsed if they felt that they had learned the layout very well. The purpose of this self-determined criterion was to prevent participants becoming bored. Seventeen out of the 30 participants moved on to the test phase before 20 minutes has passed (mean 16 min, SD 2 min). We used this subjective criterion because experimenters could check the participants’ objective memory performance afterwards and let the participants revisit the building and learn the layout again if performance was sub-optimal, before they proceeded to the scanner. Four out of the 30 participants (only one of whom was among the 17 participants who asked to move on to testing prior to the 20 minutes elapsing) had to re-visit the virtual building for up to 5 additional minutes because the accuracy either in the egocentric judgment test or the short practice for the scanning object-location test was below 70%. These four participants’ accuracy during scanning was between 78% and 83%. The mean accuracy of the 30 participants for the scanning task was 93% (SD 5.5%). We are confident, therefore, that every participant had good knowledge of the spatial layout.

#### Pre-scan egocentric judgment task

Immediately after the learning phase, there was a spatial memory test which required participants to make egocentric spatial judgments in the gallery. This test was used to examine the influence of vertical and horizontal boundaries on the mental representation of 3D space (see the behavioural analysis section).

On each trial of this test, participants saw a short dynamic video which provided the sensation of being transported to one of the 16 paintings (locations) from the corridor (duration 2.5 s; Fig. 1B, Supplementary Fig. 1). These videos were used to promote the impression of navigation whilst providing full experimental control. On half of the trials, participants started from one end of the corridor facing a floor sign on the wall, while on the remaining trials, they started from the other end of the corridor facing the stairs. In both cases they would terminate in the same location within a room, regardless of whether they began the journey facing the floor sign or stairs.

On every trial, participants were transported to one of two rooms on the floor where they started - thus the videos did not contain vertical movement via the staircase. Of note, except for the target painting, the other three paintings in a room were concealed behind curtains (Fig. 1B). Once a participant arrived at the target painting within a room, a question appeared on the screen. The question asked about the position of another painting relative to the participant’s current position, e.g. “Is the pig on your left?”, “Is the sunflower on your right?”, “Is the dog above you?”, “Is the duck below you?”. Participants responded yes or no by pressing a keypad with their index or middle finger. Similar to a previous study by Marchette et al. (2014), we instructed participants to interpret the left/right/above/below broadly, “including anything that would be on that side of the body” and not just the painting directly left/right/above/below. For example when a participant was facing the turtle painting in Fig. 1C, the sunflower was on their right and the duck was above. The time limit for answering the question was up to 5 s. The inter-trial interval (ITI) was drawn from a truncated gamma distribution (mean 2.9 s, minimum 2.0 s, maximum 6.0 s, shape parameter 4, scale parameter 0.5) and there were 64 trials. Participants were provided with their total number of correct and wrong answers at the end of the test, but did not receive feedback on individual trials.

#### fMRI scanning object location memory task

On each trial of the scanning task, participants were transported to one of the paintings from the corridor as in the pre-scan memory test (duration 2.5 s; Fig. 1B). All four paintings in the room were concealed behind curtains. Once a participant arrived at a painting, the curtain was lifted. The participant then indicated whether the painting was the correct one or not for that location by using an MR-compatible keypad. On 80% of the trials, the correct painting was presented and on 20% of trials a painting was replaced by one of the other 15 paintings. The response to the question was self-paced with an upper limit of 4.5 s (mean response time 1.3 s, SD 0.7 s), and ITIs were the same as those in the pre-scan memory test. There were 100 trials for each scanning session and each participant completed 4 scanning sessions with a short break between them, making a total functional scanning time of ˜50 minutes. Participants were told the total number of correct and wrong answers at the end of each scanning session, but individual trial feedback was not given.

The order of visiting the paintings (locations) was designed to balance first-order carry-over effects (see Aguirre 2007; Nonyane and Theobald 2007). This balancing sequence has been used in other fMRI studies involving repetition suppression (e.g., Vass and Epstein, 2013; Sulpizio et al., 2014). This meant that one location was followed by every other location with similar frequency. One issue relating to this sequence is that the trials in which the corner or room, or both, were repeated, occurred less frequently than trials where none were repeated. This is because the chance of visiting the same corner (or room) in the following trial is 25% as there are 4 possible corners (or rooms). This resulted in different numbers of trials being included in the “same corner” and “different corner” conditions (or “same room” and “different room” conditions). For maximum statistical power when contrasting these conditions, ideally every regressor would contain a similar number of trials and as many trials as possible. We nevertheless found robust and dissociable effects when we contrasted the same and different corner (or room) conditions (see Results section). Furthermore, when we shuffled the trial identity by randomly assigning the “same corner” and “different corner” labels and preserving the ratio of trials, we did not observe significant corner or room encoding in any of our ROIs, or anywhere else in the brain. This implies that a difference in the number of trials included in each condition does not invariably generate a significant difference in BOLD activity. Each participant visited the locations in a different order because we used random permutation for the association between the actual location and the index in the sequence that balanced the carry-over effects.

### Behavioural analyses

#### Pre-scan egocentric judgment test

To test the influence of compartmentalisation by vertical and horizontal boundaries on spatial judgments, we compared the accuracy and response time (RT) of egocentric spatial judgments between four conditions (Fig. 1C): (1) within; when the painting in question was in the same room as a participant; e.g. a participant is facing the turtle and made a spatial judgment about the pig, rose, or koala; (2) vertical; when the painting in question was in the room above or below a participant; e.g. a participant facing the turtle was asked about the dog, gold fish, duck or chicken; (3) horizontal; when the painting in question was in the adjacent room on the same floor as a participant; e.g. a participant facing the turtle was asked about the sunflower, crab, dolphin or swan; (4) diagonal; when the painting in question was in a diagonal room; e.g. a participant facing the turtle was asked about the elephant, eagle, lily or tulip.

If participants had a holistic mental representation of 3D space irrespective of physical boundaries within the building, performance for all four conditions should be similar. In contrast, if their mental representation was segmented into each room, spatial judgments within the same room (within) would be facilitated and therefore higher accuracy and/or faster response times would be expected compared to spatial judgments across different rooms (vertical, horizontal or diagonal conditions). If space is predominantly divided into a horizontal plane, as suggested by some previous studies (Jovalekic et al. 2011; Thibault et al. 2013; Flores-Abreu et al. 2014), spatial judgments about paintings on different floors (vertical, diagonal) would be more difficult than paintings on the same floor (within, horizontal). We used a repeated one-way ANOVA and post-hoc paired t-tests to compare the accuracy and RT for the four conditions.

#### Object location memory test during scanning

We tested whether spatial knowledge of 3D location was organised into multiple compartments by measuring a behavioural priming effect. Each trial was labelled as one of four conditions depending on the room participants visited in the immediately preceding trial (Fig. 2A, B). Figure 2B shows an example trial sequence and the room label for each trial in red: (1) same; when participants visited the same room in the previous trial, e.g. the 2^nd^ trial; (2) vertical; when participants previously visited the room above or below the current room, e.g. the 3^rd^ trial; (3) horizontal; when participants previously visited the adjacent room on the same floor, e.g. the 5^th^ trial; (4) diagonal; when participants previously visited neither vertically nor horizontally adjacent room, the e.g. 4^th^ trial. A holistic, volumetric representation of space would result in similar behavioural performance for all four conditions. If representations were compartmentalised, participants would make more accurate and/or faster judgments when spatial memory was primed by the representation of the same compartment (room). If spatial representations were further grouped along the horizontal plane, visiting the adjacent room on the same floor (horizontal condition) will also evoke a behavioural priming effect. Alternatively, the space might be represented in a vertical column, leading to a prediction of a priming effect for the vertical condition. We compared accuracy and RT for the four conditions using a repeated-measure ANOVA and post-hoc paired t-tests.

**Figure 2.**
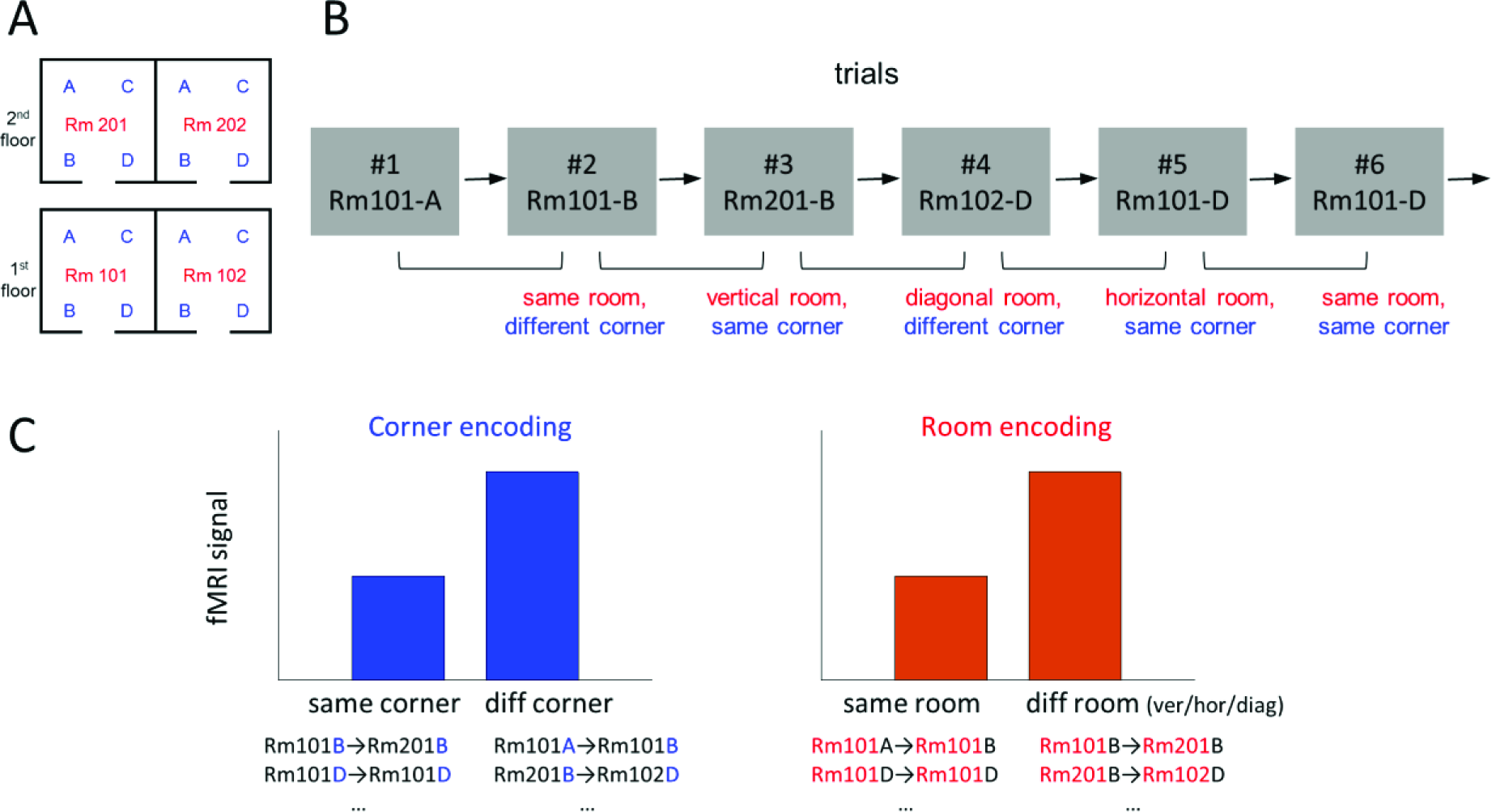
Analysis overview. (A) A floor plan of the virtual building. The 4 rooms are labelled as “Rm101”, “Rm102”, “Rm201”, “Rm202” and the 4 corners as “A”, “B”, “C”, “D” for the purposes of explanation here. Participants were not told of any explicit labels during the experiment. (B) An example trial sequence. For the behavioural and fMRI repetition suppression analyses, each trial was labelled based on its spatial relationship with the preceding trial, e.g., the 2^nd^ trial belongs to the “same room, different corner” condition. Of note, this trial definition is used for analysis only and participants were not asked to pay attention to the preceding trial. (C) Predictions for the fMRI signals. If some brain regions encode corner information, lower fMRI signal is expected for the same corner condition compared to the different corner condition. If room information is encoded, fMRI signal is expected to be lower for the same room condition compared to the different room condition.

### Scanning and pre-processing

T2*-weighted echo planar images (EPI) were acquired using a 3T Siemens Trio scanner (Siemens, Erlangen, Germany) with a 32-channel head coil. Scanning parameters optimised for reducing susceptibility-induced signal loss in areas near the orbitofrontal cortex and medial temporal lobe were used: 44 transversal slices angled at -30°, TR=3.08 s, TE=30 ms, resolution=3×3x3mm, matrix size=64x74, z-shim gradient moment of -0.4mT/m ms (Weiskopf et al. 2006). Fieldmaps were acquired with a standard manufacturer’s double echo gradient echo field map sequence (short TE=10 ms, long TE=12.46 ms, 64 axial slices with 2 mm thickness and 1 mm gap yielding whole brain coverage; in-plane resolution 3 x 3 mm). After the functional scans, a 3D MDEFT structural scan was obtained with 1mm isotropic resolution.

Preprocessing of data was accomplished using SPM12 (www.fil.ion.ucl.ac.uk/spm). The first 5 volumes from each functional session were discarded to allow for T1 equilibration effects. The remaining functional images were realigned to the first volume of each run and geometric distortion was corrected by the SPM unwarp function using the fieldmaps. Each participant’s anatomical image was then coregistered to the distortion corrected mean functional images. Functional images were normalised to MNI space, then spatial smoothing (FWHM=8mm) was applied.

### fMRI analyses

#### Main analysis: room and corner encoding

We used an fMRI repetition suppression analysis to search for two types of spatial information in the brain: (1) *corner*; a participant’s location within a room, and (2) *room*; which room a participant was in. fMRI repetition suppression analysis is based on the assumption that when a similar neural population is activated across two consecutive trials, the fMRI signal is reduced during the second trial. Therefore, if a brain region encodes the corner information, visiting the same corner in a consecutive trial would result in reduced fMRI signal compared to visiting a different corner (Fig. 2C). For example, visiting Rm201-B after Rm101-B (3^rd^ trial in the example sequence, Fig. 2B) or visiting Rm101-D twice in a row (6^th^ trial in the example) would evoke reduced fMRI signal than visiting Rm101-B after Rm101-A (2^nd^ trial in the example) or visiting Rm102-D after Rm201-B (4^th^ trial). On the other hand, if a brain region encodes room information, visiting the same room in consecutive trials (e.g. in Fig. 2B, Rm101-A → Rm101-B, 2^nd^ trial) would result in reduced fMRI signal compared to visiting a different room (e.g. Rm101-B →Rm201-B, 3^rd^ trial).

We also tested whether vertical and horizontal boundaries similarly influenced neural similarity between rooms. If there was a bias in encoding horizontal information better than vertical (floor), then the two rooms on top of each other (e.g. Fig. 2B, Rm101 and Rm201) would be less distinguishable than the two rooms on the same floor (e.g. Rm101 and Rm102). Therefore, visiting a vertically adjacent room (e.g. Rm101-B → Rm201-B, 3^rd^ trial) would result in more repetition suppression, leading to a reduced fMRI signal, than visiting a horizontally adjacent room (e.g. Rm102-D → Rm101-D, 5^th^ trial). We were also able to ask whether two rooms in a diagonal relationship (e.g. Rm201-B → Rm102-D, 4^th^ trial) were more distinguishable than vertically or horizontally adjacent rooms.

To answer these questions, we constructed a GLM which modelled each trial based on its spatial relationship to the preceding trial in terms of two factors: corner and room. The corner factor had 2 levels: same or different corner, and the room factor had 4 levels: same, vertical, horizontal or diagonal room. This resulted in a 2 x 4 = 8 main regressors. Each regressor was a boxcar function which for each trial modelled the entire stimulus duration including the virtual navigation period (2.5 s) and subsequent object-location memory test (mean RT 1.3 s, SD 0.7 s) (Fig. 1B top three panels) convolved with the SPM canonical hemodynamic response function. Information about spatial location was cumulatively processed throughout the navigation video and continued until participants decided whether the painting was the correct one or not for the location, and hence we modelled the entire period as a single boxcar. Moreover, this fMRI study was not designed to distinguish between temporally adjacent events or cognitive processes, because we favoured a more naturalistic task (i.e., without the delays that jitter would introduce). The first trial of each scanning session, which did not have an immediately preceding trial, or the trials where participants were incorrect (mean 6.8%, SD 5.5%) were excluded from the main regressors and modelled separately. The GLM also included nuisance regressors: six head motion realignment parameters and the scanning session-specific constant regressor.

First, we conducted a whole-brain analysis to search for corner and room information using two contrasts: (1) “same corner < different corner”, collapsed across the room factor, and (2) “same room < different room (=average of the vertical room/horizontal room/diagonal rooms)”, collapsed across the corner factor. Each participant’s contrast map was then fed into a group level random effects analysis. Given our a priori hypothesis about the role of hippocampus and retrosplenial cortex (RSC) for encoding spatial information, we report voxel-wise p-values corrected for anatomically defined hippocampus and RSC regions-of-interest (ROIs). For the rest of the brain, we report regions that survived a whole-brain corrected family-wise error (FWE) rate of 0.05.

The hippocampus and retrosplenial cortex ROIs were manually delineated on the group-averaged structural MRI scans from a previous independent study on 3D space representation (Kim et al., 2017) (Supplementary Figure 3). The bilateral hippocampus mask contained the whole hippocampus from head to tail (304 voxels in 3 x 3 x 3 mm EPI resolution). Although some navigation fMRI studies have defined a “retrosplenial complex” which includes Brodmann areas 29-30, occipitotemporal sulcus and posterior cingulate cortex (e.g., Marchette et al. 2014), we used a more precise anatomical definition of RSC, based on cytoarchitecture, which includes only Brodmann areas 29-30 (Vann et al. 2009) (293 voxels in 3 x 3 x 3 mm EPI resolution). Functionally defined ROIs vary from one study to another depending on the statistical threshold and individual differences. We would expect that the anatomical RSC and functionally defined RSC overlap, and whether an anatomical or functional definition is more appropriate depends on the specific research question. In our study, we were interested in finding corner or room information across the entire brain, with precise anatomical priors in hippocampus and RSC, and so we preferred a conservative and threshold-free anatomical definition.

Having identified brain regions that contained significant corner information from the whole brain analysis, we examined the spatial encoding in these regions further by extracting the mean fMRI activity. As a proxy for the mean fMRI activity, beta weights for every voxel within the spherical ROIs (radius 5mm, centred at the peak voxel) were averaged for each participant, and then compared at the group level by paired t-tests. For this functional ROI-based analysis, we divided the “same corner” condition into “same corner, same room” and “same corner, different room” and compared each condition to “different corner”. This analysis allowed us to rule out the possibility that the corner encoding was driven purely by the repetition suppression effect of “same corner, same room” < “different corner”. If a brain region encodes each of the 16 locations (or associated paintings) without a spatial hierarchy, repetition suppression would only occur for the “same corner, same room” condition and there would be no difference between “same corner, different room” and “different corner”.

We conducted a similar control analysis in the brain regions that contained significant room information (“same room < different room”). We compared the mean activity of “same room, same corner” and “same room, different corner” to “different room” to rule out the possibility that the room encoding was driven by the repetition suppression of the exactly same location. Crucially, we also compared mean activity of different room conditions (vertical/horizontal/diagonal rooms) to test for any potential bias in encoding vertical or horizontal information.

Note that our experiment was not designed to test a pure non-hierarchical encoding model. While the contrast “same location < all others” can be used to identify brain regions expressing non-hierarchical encoding, this contrast is not orthogonal to the main effect contrasts (“same corner< different corner” or “same room<different room”). Moreover, this analysis is subject to the criticism that any brain areas identified could be responding to the identity of the paintings rather than the 3D locations, given that each location was associated with unique painting. Our study was specifically designed to examine the main effects of corner and room information. We therefore used straightforward classical statistics (t-tests) to determine these main effects, and then interrogated these findings further to establish that the main effects were not solely driven by the repetition suppression for the same location.

The data could also be examined in terms of 3D physical metric distance from the preceding trial modelled as a linear parametric regressor. However, the highly discretized nature of the environment makes inferences about metric encoding difficult in this context, and this issue would be better addressed with a different type of environment.

#### Supplementary analysis: room versus view encoding

In this experiment, room information was cued by a distinctive view such as a wall containing a floor sign, therefore the room encoding effect could arise due to view encoding and/or more abstract spatial information about a room that was not limited to a particular view. We were able to test for these possibilities because participants were virtually transported to each room from two directions (Supplementary Fig. 1), which means they could visit the same room on consecutive trials from the same or different direction. For example, if they had visited Rm101 from the floor sign side in the preceding trial and visited Rm101 from the stair side in the current trial, the views were very different even though the same room was visited. On the other hand, if they had visited Rm101 from the floor sign side in the preceding trial and visited Rm201 from the floor sign side in the current trial, the views were similar even though two rooms were different.

We constructed a GLM which modelled each trial based on two factors: whether it was the same or a different room from the previous trial, and whether the starting direction (view) was the same or different direction from the previous trial. This resulted in 4 trial types: “same room, same view”, “same room, different view”, “different room, similar view”, and “different room, different view”. As in the main analysis, only correct trials were included for the main regressors, and head motion realignment parameters and scanning session-specific constant regressors were included in the GLM. For each participant, we extracted the mean activity (beta weights) for each trial type in the room encoding regions identified in the “same room < different room” contrast described earlier. We conducted a repeated measures ANOVA and post-hoc paired t-tests to compare the mean beta weights between the “same room, same view”, “same room, different view”, and “different room” (collapsed over similar and different view). If only the view was encoded, then the “same room, same view” would have a reduced fMRI signal compared to “different room”, but “same room, different view” would not be associated with a reduced fMRI signal compared to the “different room” condition. If abstract room information was encoded, the “same room, different view” condition would also be associated with reduced fMRI signal compared to the “different room” due to repetition of the room. We were also able to compare “same room, same view” and “same room, different view” to test view dependency when the room was repeated.

## Results

### Behavioural results

#### Pre-scan egocentric judgment task

In order to examine the influence of vertical and horizontal boundaries on the mental representation of 3D space, we compared the accuracy and RT of spatial judgments for 4 conditions: within, vertical, horizontal and diagonal rooms. Participants were faster at judging the location of paintings within the same room compared to paintings in different rooms (Fig. 1D; F(3,87)=5.4, p=0.002, post-hoc paired t-tests: within vs. vertical, t(29)=-3.5, p=0.001; within vs. horizontal, t(29)=-2.7, p=0.011; within vs. diagonal, t(29)=-2.2, p=0.034). There was no significant difference in RT between the vertical, horizontal and diagonal rooms. This result suggests the importance of a physical boundary, but this was not influenced by whether the boundary was vertical or horizontal. Accuracy did not differ significantly between the four conditions (F(3,87)=1.2, p=0.3; mean overall accuracy 80%, SD 12%).

#### Object-location memory task during scanning

Overall, participants performed well on the object-location memory task (mean accuracy 93%, SD 5.5%). Participants were more accurate and faster at judging whether a painting was in the correct location if they had visited the same room in the preceding trial (Fig. 3A, B; Accuracy: F(3,87)=4.2, p=0.008; post-hoc paired t-tests: same vs. vertical, t(29)=3.4, p=0.002; same vs. horizontal, t(29)=2.3, p=0.032; same vs. diagonal, t(29)=2.3, p=0.028; RT: F(3,87)=8.3, p<0.001; same vs. vertical, t(29)=-4.0, p<0.001; same vs. horizontal, t(29)=-2.8, p=0.009; same vs. diagonal, t(29)=-4.1, p<0.001), and neither accuracy nor RT differed between the vertical, horizontal or diagonal rooms. This result, along with the pre-scan memory task, suggests the mental representation of 3D space was segmented into each room, regardless of vertical floor.

**Figure 3.**
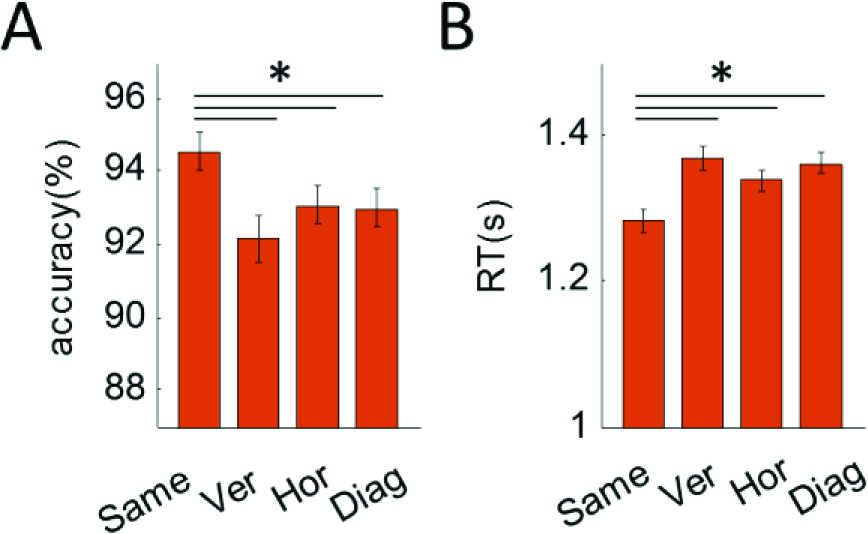
The behavioural priming effect of room during the scanning task. (A) Accuracy was significantly higher for the same room condition compared to all other rooms. There was no significant difference between the different room types. (B) RT was significantly reduced for the same room condition compared to all other conditions. There was no significant difference between different room types. Error bars are SEM adjusted for a within-subjects design (Morey 2008). *p<0.05.

### fMRI results

We tested if the brain represents a multi-compartment 3D building space in a hierarchical manner by separately encoding the corner (“where am I within a room?”) and room (“in which room am I in the building?”). We searched for these two types of information using an fMRI repetition suppression analysis. Furthermore, we investigated whether there were differences in how vertical and horizontal information was encoded. We present the results for our two ROIs – the RSC and hippocampus – and any other region that survived whole brain correction – there was only one, the right parahippocampal cortex.

#### Corner information

The “same corner < different corner” contrast revealed left anterior lateral hippocampus (Fig. 4, peak MNI coordinate [-33,-19,-16], t(29)=5.31, p=0.001, small volume corrected with a bilateral hippocampal mask), suggesting that this region encodes which corner a participant is located within a room. No other brain region showed a significant corner repetition suppression effect at the whole brain corrected level.

**Figure 4.**
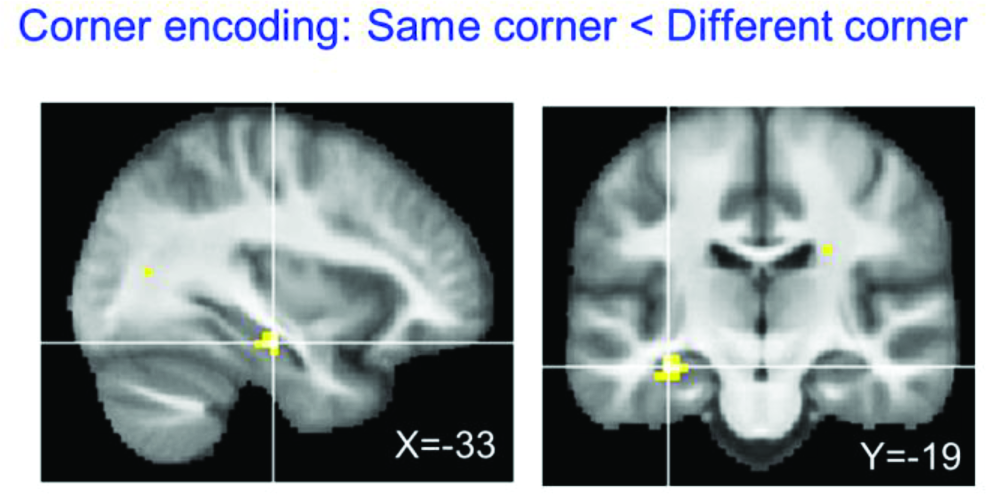
Corner encoding regions. The whole brain contrast “same corner < different corner” revealed only the left anterior hippocampus (peak MNI=[-33, -19, -16], t(29)=5.31, p<0.001). The thresholded map is overlaid on the group average structural MRI scan (p<0.001, uncorrected for display purposes). No other brain region survived multiple comparison correction.

We further examined the spatial encoding in the left anterior lateral hippocampus by extracting the mean activity (beta) for each condition. We investigated the fMRI signal when exactly the same location was visited (“same corner, same room”, e.g. Fig. 2B, Rm101-A → Rm101-A) and when the same corner, but a different room was visited (“same corner, different room”, e.g. Rm101-A → Rm201-A) and compared them to the “different corner” condition (e.g. Rm101-A → Rm201-B). If the entire building is represented in a single volumetric space without a hierarchy, then each of the locations would be uniquely encoded, so repetition suppression is expected only for the “same corner, same room” condition. Our finding speaks against the single volumetric representation hypothesis because both “same corner, same room” and “same corner, different room” conditions evoked significant repetition suppression effects compared to the “different corner” condition (one-sided paired t-tests: “same corner, same room” < “different corner”, t(29)=-4.4, p<0.001; “same corner, different room” < “different corner”, t(29)=-4.2, p<0.001). This implies that the anterior hippocampus contains local corner information that is generalised across different rooms, supporting an efficient hierarchical representation of 3D space.

On a related note, one might ask whether the hippocampus showed sensitivity to the heading direction instead of location. Participants faced opposite walls when they were at corner A (or B) and when they were at corner C (or D) (Figure 1A, 2A). However, they faced the same direction when they were at location A and B (or C and D) and further analysis revealed that there was no difference in the anterior hippocampus when participants visited a corner on the same wall or the opposite wall. Thus, we can conclude that the hippocampus encoded corner information rather than heading direction.

#### Room information

The “same room < different room” contrast revealed bilateral RSC (right RSC peak [9,-52,11], t(29)=8.55, p<0.001; left RSC peak [-9, -58, 14], t(29)=7.91, p<0.001, small volume corrected with a bilateral RSC mask), right parahippocampal cortex (peak [27, -37,-16], t(29)=7.21, p<0.001), and the posterior part of the hippocampus (right hippocampus peak [27, -28, -10], t(29)=6.12, p<0.001; left hippocampus peak [-27,-34,-7], t(29)=4.32, p=0.014, small volume corrected with a bilateral hippocampus mask) (Fig. 5A). This suggests that these regions encode in which room a participant was located in the building. It is notable that the room information was detectable in the posterior portion of hippocampus, compared to corner information which was detectable in the anterior hippocampus.

**Figure 5.**
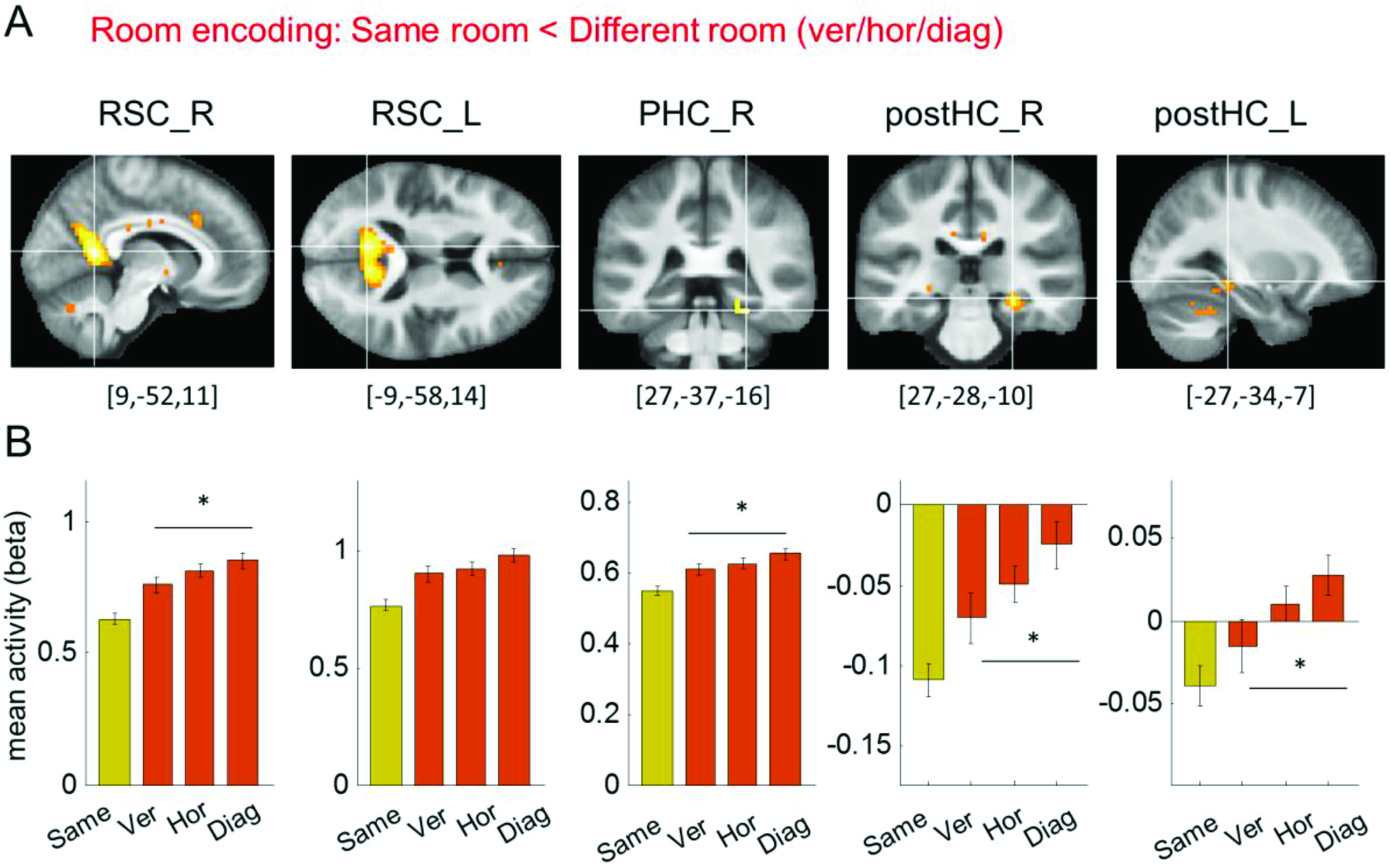
Room encoding regions. (A) The whole brain contrast “same room < different room” revealed bilateral RSC (RSC_R, RSC_L), right parahippocampal cortex (PHC_R) and bilateral posterior HC (postHC_R, postHC_L). Given our a priori interest in RSC and posterior hippocampus, their clusters are shown with a small volume corrected threshold level (t(29)>3.67, t(29)>3.75), while the parahippocampal cortex cluster is shown with a whole-brain corrected threshold (t(29)>6.01). The peak MNI coordinate is shown below each cluster. (B) Comparison of mean activity for three different room types (vertical/horizontal/diagonal) at each cluster (5mm sphere at peak voxel). The “same” condition (in yellow) is shown for reference purposes. The response to the diagonal condition was significantly larger than for the vertical condition in all regions except the left RSC. There was no significant difference between the vertical and horizontal conditions. Error bars are SEM adjusted for a within-subjects design (Morey 2008). *p<0.05.

We further examined spatial encoding in the right and left RSC (RSC_R, RSC_L), right parahippocampal cortex (PHC_R) and right and left posterior hippocampus (postHC_R, postHC_L) by extracting the mean fMRI activity for the “same corner, same room”, “different corner, same room”, and “different room” conditions. In all regions, we found significant repetition suppression effects for both “same corner, same room” and “different corner, same room” conditions compared to the “different room” (one-sided paired t-tests: “same corner, same room” < “different room”: RSC_R, t(29)=-6.6, p<0.001; RSC_L, t(29)=-4.9, p<0.001; PHC_R, t(29)=-4.9, p<0.001; postHC_R, t(29)=-4.9, p<0.001; postHC_L, t(29)=-3.2, p=0.002; “different corner, same room” < “different room”: RSC_R, t(29)=-4.1, p<0.001; RSC_L, t(29)=-3.6, p<0.001; PHC_R, t(29)=-5.0, p<0.001; postHC_R, t(29)=-3.0, p=0.003; postHC_L, t(29)=-2.2, p=0.02). These findings suggest the presence of room information that is independent of the local corner.

We then tested for the existence of vertical-horizontal asymmetry in these five room encoding regions - RSC_R, RSC_L, PHC_R, postHC_R, postHC_L - by extracting the mean activity for sub-categories of the different room conditions: vertical room, horizontal room and diagonal room (Fig. 5B). If vertical information was relatively poorly encoded compared to horizontal information, we would expect that two rooms on top of each other (e.g. Rm101 and Rm201, Fig. 2A) to be more similarly represented in the brain than the two adjacent rooms on the same floor (e.g. Rm101 and Rm 102). Consequently, we would expect less fMRI activity for the vertical room condition than the horizontal condition. We also tested whether two rooms in a diagonal relationship were more distinguishable than either the vertically or horizontally adjacent room due to physical or perceptual distance. For this comparison, we used a repeated measures ANOVA with 3 room types as a main factor. In Fig. 5B, we also plot the same room condition for reference purposes. Since the room encoding region was defined by the “same room < different room” contrast, the “same room” should be associated with reduced activity in all regions. We found a significant main effect in all regions except for the left RSC (RSC_R, F(2,58)=3.8, p=0.029; RSC_L, F(2,58)=2.4, p=0.10; PHC_R, F(2,58)=3.2, p=0.049; postHC_R, F(2,58)=3.5, p=0.036; postHC_L, F(2,58)=3.6, p=0.032). Post-hoc t-tests showed that this main effect was driven by a small difference between the vertical and diagonal conditions (“ver” versus “diag”, RSC_R, t(29)=-2.4, p=0.022; PHC_R, t(29)=-2.5, p=0.017; postHC_R, t(29)=-2.3, p=0.031; postHC_L, t(29)=-2.3, p=0.028). The diagonal condition evoked a larger signal than the vertical condition, implying that two rooms in a diagonal relationship are more differently encoded than two rooms on top of each other. None of the regions showed a significant difference between the vertical and horizontal conditions.

As a side note, the sign of the mean activity (beta) was negative in the hippocampus, implying that the activity was lower during the stimulus presentation period (virtual navigation and subsequent object-location memory test) compared to the fixation cross inter-trial-interval (ITI). In the literature the hippocampus is often reported to show negative beta values during stimulus presentation or task periods (e.g. Bakker et al. 2008; Evensmoen et al. 2015; Hodgetts et al. 2015; Brodt et al. 2016). We believe that the absolute beta value of a single condition has little meaning in our study as the implicit baseline (ITI period) was not a meaningful experimental condition. Our study explicitly focussed on comparisons between the main experimental conditions such as the “same room” versus the “horizontal room”. The comparisons showed the predicted pattern of repetition suppression, with the fMRI signal associated with the “same” condition reduced compared to the different room conditions.

#### Supplementary analysis: room versus view encoding

In order to know whether the RSC, parahippocampal cortex and posterior hippocampus encoded view information associated with each room and/or more abstract spatial knowledge about the room, we conducted a supplementary analysis that separated the same room condition into sub-categories of same view and different view conditions. We then compared them to the different room condition (see Methods and Supplementary Fig. 1). We observed repetition suppression effects even when participants visited the same room but approached it from a different view (Supplementary Fig. 2; one-sided paired t-tests: “same room, different view” < “different room”, RSC_R, t(29)=-2.1, p=0.021; RSC_L, t(29)=-2.4, p=0.011; PHC_R, t(29)=-1.9, p=0.034; postHC_R, t(29)=-1.8, p=0.041; postHC_L, t(29)=-1.9, p=0.034). This suggests that these regions contained abstract room information that was not limited to the exact view. However, there was also evidence for view encoding in some regions. For example, visiting the same room from the same view evoked significantly less activity compared to visiting the same room from different view in the right RSC, PHC_R and the postHC_R (one-sided paired t-tests: “same room, same view” < “same room, different view”, RSC_R, t(29)=-2.5, p=0.010; PHC_R, t(29)=-3.4, p=0.001; postHC_R, t(29)=-1.8, p=0.041). In contrast, the left RSC and left posterior hippocampus did not show any significant differences between the same view and different view (p>0.1). In summary, left RSC and left posterior hippocampus showed relatively pure room encoding that was independent of view. Other regions showed additional view dependency, and this was particularly strong in right parahippocampal cortex.

## Discussion

In this study we investigated how a multi-compartment 3D space (a multi-level gallery building) was represented in the human brain using behavioural testing and fMRI repetition suppression analyses. Behaviourally, we observed faster within-room egocentric spatial judgments and a priming effect of visiting the same room in an object-location memory test, suggesting a segmented mental representation of space. At the neural level, we found evidence of hierarchical encoding of this 3D spatial information, with the left anterior lateral hippocampus containing local corner information within a room, whereas RSC, parahippocampal cortex and posterior hippocampus contained information about the rooms within the building. Furthermore, both behavioural and fMRI data were concordant with unbiased encoding of vertical and horizontal information.

We consider first our behavioural findings. There is an extensive psychological literature suggesting that space is encoded in multiple “sub-maps” instead of a flat single map. Accuracy and/or reaction time costs for between-region spatial judgments (McNamara et al. 1989; Montello and Pick 1993; Han and Becker 2014), and context swap errors, where only the local coordinate is correctly retrieved (Marchette et al. 2017), are evidence for multiple or recurring sub-maps. Here, we observed faster RT for within-room direction judgments and a behavioural priming effect of visiting the same room during a spatial memory task. Our findings are therefore consistent with the idea of a segmented representation of space.

Importantly, the current study examined regionalisation in 3D space and compared the influence of vertical and horizontal boundaries for the first time. Some previous studies have suggested a bias in dividing space in the horizontal plane (Jovalekic et al. 2011; Thibault et al. 2013; Flores-Abreu et al. 2014). The horizontal planar encoding hypothesis predicts an additional behavioural cost for spatial judgments across floors and priming effects for the rooms within a same floor. However, we did not find any significant difference in performance for spatial judgments across vertical and horizontal boundaries, or priming effects for rooms on the same floor. Although the absence of significant difference does not necessarily mean equivalence, the most parsimonious interpretation would be that each room within our 3D space was similarly distinguishable.

This fits with the symmetric encoding of 3D location information in a semi-volumetric space previously reported in bats and humans (Yartsev and Ulanovsky 2013; Kim et al. 2017). One concern might be that the small number of rooms in our virtual building allowed participants to encode each room categorically without being truly integrated in a 3D spatial context. However, in order to be successful at the egocentric judgments task across rooms (mean accuracy was 80%), our participants must have had an accurate representation of the 3D building. Testing an environment with more floors and rooms in the future could facilitate the search for any additional hierarchies within 3D spatial representations. For example, rooms might be further grouped into the horizontal plane or a vertical column in a more complex environment. It might also help to reveal subtle differences, if they exist, between vertical and horizontal planes.

Considering next our fMRI results, we found that fMRI responses in the left anterior lateral hippocampus were associated with local corner information that was generalised across multiple rooms. This fits well with previous findings that hippocampal place cells in rodents fire at similar locations within each segment of a multi-compartment environment (Derdikman et al. 2009; Spiers et al. 2015). This common neural code enables efficient encoding of information. For example, the 16 locations in our virtual building could be encoded using only 8 unique codes (4 for distinguishing the corners of rooms and 4 for distinguishing the rooms themselves) given its regular substructures. This room-independent representation in the anterior lateral hippocampus can also be seen as a ‘schematic’ representation of space (Marchette et al. 2017) where the regular structure of the environment is extracted. Furthermore, there is evidence that the ability of the hippocampus to extract regularity in the world is not limited to the spatial domain. A previous fMRI study found that temporal order information in the hippocampus generalised across different sequences (Hsieh et al. 2014). Statistical learning of temporal community structure has also been associated with the hippocampus (Schapiro et al. 2016) and, interestingly, localised to the anterior portion. Rodent electrophysiology and modelling work also suggests that ventral hippocampus (analogous to the human anterior hippocampus) is well suited to generalising across space and memory compared to dorsal hippocampus (analogous to the human posterior hippocampus) (Keinath et al. 2014).

In addition to generalised within-room information, it is also important to know a room’s location to identify one’s exact position within a building. We found that multiple brain regions represented room information, with the RSC exhibiting the most reliable room repetition effect. At first this finding might seem surprising, given that head direction information has been consistently associated with the RSC in humans and rodents (Baumann and Mattingley 2010; Marchette et al. 2014; Jacob et al. 2016; Shine et al. 2016). In our virtual building, participants faced paintings on opposite walls within a room. Therefore, if RSC encoded the participant’s facing direction, local corner encoding would be expected instead of room encoding. However, numerous findings suggest that RSC encodes more than head direction; processing of multiple spatial features such as location, view, velocity and distance have been linked with this region (Cho and Sharp 2001; Sulpizio et al. 2014; Alexander and Nitz 2015; Chrastil et al. 2015). Moreover, RSC was found to be involved in both a location and an orientation retrieval task when participants viewed static pictures of an environment during fMRI (Epstein et al. 2007). Given the rich repertoire of spatial, visual and motor information the RSC processes, it is perhaps not surprising that some studies observed local head direction signals and others found global head direction information in this region (Marchette et al. 2014; Shine et al. 2016). This might also be influenced by functional differences within the RSC, or indeed laterality effects. In our experiment, the right RSC showed stronger repetition suppression when participants visited the same room from same view compared to when they visited the same room from different view, whereas the left RSC’s response was only influenced by the repetition of the room.

RSC might have a role in integrating local representations within a global environment. A recent theory about the neural encoding of large-scale 3D space proposed that 3D space is represented by multiple 2D fragments, and RSC is a candidate area for stitching these together (Jeffery et al. 2015). The authors’ argument was based on the reasoning that the RSC is suitable for updating orientation in multiple adjoining, sloped planes. In our experiment, room information can be broadly viewed as the orienting cue within a building that allows integration of the fragmented space. For localisation and orientation of local representations within a larger spatial context, landmark information is crucial. In the current experiment, room information was cued by salient landmarks such as the floor sign or the staircase. Landmark information could, therefore, be the key to understanding the RSC’s various spatial functions including the representation of abstract room information, scene perception, processing of directional signals and the integration of multiple local reference frames. RSC is known to support the learning of and processing of stable landmarks (Auger et al. 2012, 2015), and its head direction signal is dominated by local landmarks (Jacob et al. 2016).

The second region that represented room information was the parahippocampal cortex. It also showed a strong view dependency in addition to room information. This contrasts with the left RSC which only showed a room repetition effect. Together these findings are consistent with the proposed complementary roles of the parahippocampal cortex and RSC in scene perception, whereby the former seems to respond in a view-dependent manner whereas the RSC represents integrative and more abstract scene information. For example, it has been shown that when participants saw identical or slightly different snapshot views from one panoramic scene, RSC showed fMRI repetition effects for both identical and different views, but parahippocampal cortex only exhibited repetition suppression for the identical view (Park and Chun 2009). In addition, multivoxel patterns in RSC have been observed to be consistent across different views from each location, whereas this was not the case for the parahippocampal cortex (Vass and Epstein 2013).

Along with RSC and parahippocampal cortex, the final area to represent room information was the posterior hippocampus. The similarity in spatial encoding between these regions might be predicted from their close functional and anatomical connectivity (Kobayashi and Amaral 2003; Kahn et al. 2008; Blessing et al. 2016). It is notable that in our previous fMRI study that examined 3D spatial representation, we also found that posterior hippocampus and RSC encoded the same type of spatial information (vertical direction) while anterior hippocampus encoded a different type of spatial information (3D location) (Kim et al. 2017). In that study, different vertical directions resulted in more distinguishable views, although direction information observed in the multivoxel patterns remained significant after controlling for low level visual similarities. Our current results do not fit precisely with accounts that associate the posterior hippocampus with a fine-grained spatial map (Poppenk et al. 2013; Evensmoen et al. 2015). In fact, our findings could be interpreted as evidence in the opposite direction, namely that coarser-grained representations of the whole building engage the posterior hippocampus. Nevertheless, overall our anterior and posterior hippocampal findings provide further evidence of functional differentiation down the long axis of the hippocampus (Baumann and Mattingley 2013; Poppenk et al. 2013; Strange et al. 2014; Zeidman and Maguire 2016).

Finally, as with our behavioural data, we also examined the fMRI data for possible differences between the horizontal and vertical planes. We did not find significant differences in fMRI amplitude between the vertical and horizontal conditions in the brain structures that contained room information. This neural finding is consistent with our behavioural results of similar accuracy and RT for spatial judgments across vertical and horizontal rooms, and similar priming effects for each room. These results fit well with an isotropic representation of 3D space, similar to our previous experiment (Kim et al. 2017).

Again, as with the behavioural data, one concern might be that each room is represented in RSC, parahippocampal cortex and posterior hippocampus in a categorical, semantic manner without consideration of their physical 3D location in building. However, as we discussed earlier, egocentric spatial judgments in the pre-scan task prevented participants from separately encoding each room without the 3D spatial context. Furthermore, we found that visiting a diagonal room evoked a larger fMRI signal than visiting a vertical room, and this finding cannot be explained if each room was encoded in a flat manner without a spatial organisation. This implies that the neural representation of two rooms in a diagonal relationship were more distinguishable than two rooms on top of each other. This might be due to the change in two axes for the diagonal room (vertical and horizontal) compared to change along only one axis for the vertical room, or simply because of a longer distance between two rooms in diagonal relationship. Distance encoding has been previously reported in parahippocampal cortex and RSC (Marchette et al. 2014; Sulpizio et al. 2014).

To disambiguate these possibilities, a larger environment consisting of multiple vertical and horizontal sections should be tested. For example, if the physical distance between the rooms is the main factor for neural dissimilarity, two rooms on the same floor that were separated by 5 other rooms (e.g. Rm101 and Rm106) would be more distinguishable than two rooms that are both vertically and horizontally adjacent (e.g. Rm101 and Rm 202, Fig. 2A). If the change in both vertical and horizontal axes always has a greater effect than the change in one axis, the diagonal rooms would be more distinguishable than horizontally or vertically aligned rooms regardless of distance. Use of a larger environment would also widen the scope for detecting subtle differences, if any, in the vertical and horizontal axes.

In addition to absolute physical distance, path or navigation distance is also a consideration. For example, a typical multi-level building like the one used in the current study has limited access points to movement across the floors. People cannot directly move up to the room above through the ceiling; rather they have to use stairs or elevators which are often sparsely located in the building. Thus, two rooms on top of each other are further apart in terms of actual navigation than two rooms side by side on the same floor, even when absolute distances are identical or the vertical rooms have even shorter physical distance than the horizontal rooms. Representation of space in the hippocampus is not only influenced by absolute distance but also by path distance (Howard et al. 2014), and it has also been suggested that topology instead of physical geometry is encoded in hippocampal place cells (Dabaghian et al. 2014). It would be intriguing to systematically investigate the effect of physical and path distance, and the potential interaction with vertical/horizontal boundaries, in future studies.

In summary, here we presented novel evidence showing that a multi-compartment 3D space was represented in a hierarchical manner in the human brain, where within-room corner information was encoded by the anterior lateral hippocampus and room (within the building) information was encoded by RSC, parahippocampal cortex and posterior hippocampus. Moreover, our behavioural and neural findings showed equivalence of encoding for vertical and horizontal information, suggesting an isotropic representation of 3D space even in the context of multiple spatial compartments. Despite multi-level environments being common settings for much of human behaviour, little is known about how they are represented in the brain. These findings therefore provide a much-needed starting point for understanding how a crucial and ubiquitous behaviour – navigation in buildings with numerous levels and rooms – is supported by the human brain.

The authors declare no competing financial interests.

